# Population comparisons reveal more social interactions are correlated with larger brains in the Lake Tanganyikan cichlid *Neolamprologus brevis*

**DOI:** 10.1101/2024.12.17.627543

**Authors:** Bin Ma, Boyd Dunster, Etienne Lein, Boshan Zhu, Weiwei Li, Zoë Goverts, Alex Jordan

## Abstract

The Social Brain Hypothesis (SBH) proposes that complex social environments drive the evolution of larger brains and specific neuroanatomical adaptations. This relationship can be difficult to study in the wild, because species that differ in social organization may also diverge in morphology, ecology, phylogeny, and other life history parameters. Here we use two populations of the shell-dwelling cichlid *Neolamprologus brevis* with contrasting social environments to test whether increased social complexity is associated with larger brain sizes or specific regional adaptations. Behavioral observations revealed similarly low feeding rates in both populations, but significantly more frequent social interaction frequencies in the population one of the populations. This population had larger total brain volumes relative to body size, with a disproportionately larger telencephalon and a smaller hypothalamus, suggesting region-specific adaptations to social demands. By integrating behavioral quantification and neuroanatomical analysis, our study highlights the importance of sociality as a driver of brain evolution and demonstrates the utility of cichlid fish as a model for testing the SBH in non-mammalian systems. These findings provide empirical support for the SBH and underscore the value of combining behavioral and morphological data in evolutionary neuroethology.

## Introduction

The evolution of brain size across animal taxa has captivated scientific attention, with diverse hypotheses proposed to explain variations in brain size and structure. Among these, the Social Brain Hypothesis (SBH) posits that species in complex social environments evolve larger brains to meet the cognitive demands of social interactions (Dunbar, 2009). While many studies support this theory, showing a correlation between brain size and social group size in primates (Dunbar, 2009), the hypothesis remains contentious. Studies on carnivores (Finarelli & Flynn, 2009) and birds (Hardie & Cooney, 2023) have yielded conflicting results, with some suggesting that ecological and developmental factors may overshadow social influences on brain size. Although large-scale phylogenetic comparisons between species are crucial for revealing genetic adaptations on evolutionary timescales, studying divergent ecomorphs can provide a higher resolution by reducing confounding factors such as genomic and morphological variation.

Although the SBH was developed to explain brain evolution in primates, this application of this hypothesis has extended to multiple taxa. Fish can be a particularly useful group in such studies due to their high brain plasticity, allowing researchers to investigate the effects of social and ecological factors on neuroanatomy over shorter timescales compared to mammals (K. Kotrschal et al., 1998). Recent study have highlighted the influence of social environments on fish brain size, with evidence suggesting that higher population densities can lead to increased brain sizes and enhanced cognitive abilities (Triki et al., 2019). However, most studies have relied on indirect proxies for social complexity, such as population density and mating systems, rather than direct behavioural measurements. Even fewer have linked observed social behaviours to neuroanatomical variations under natural conditions, leaving the connection between social interactions and brain morphology largely unexplored.

To address this gap, our study investigates two divergent populations of the shell-dwelling cichlid *Neolamprologus brevis* that differ markedly in social complexity. In populations at Chezi this species lives on sandy bottoms in sparsely scattered shells where they form pairs inhabiting a single shell, In Chikonde this species inhabits continuous shell beds alongside many other heterospecifics, and each individual inhabits a separate shell. While there is some debate about the species status of these populations, an exhaustive study (Ota et al., 2012) concluded these different habitats represent divergent ecomorphs (and potentially alternative reproductive tactics) of the single species *N. brevis*. The ecological environments of these forms, characterized by similar ecotypes, comparable predation pressures and food availability, offer a rare opportunity to control confounding ecological factors and focus on the influence of social structure. By quantifying specific behaviours (social and foraging) and analysing brain size and structure, this study aims to test whether increased social complexity is associated with larger overall brain size and specific neuroanatomical adaptations.

We hypothesize that the population of *N. brevis* living in continuous shell beds will exhibit more frequent social interactions, have larger total brain volumes. Additionally, we expect relative enlargement of brain regions associated with social information processing.

## Methods

### 1. Study Species and Ecology

We studied two forms of the shell-dwelling cichlid *Neolamprologus brevis* from Lake Tanganyika that differ in social organization but share similar ecological environments (**Figure 1**).

**Figure 1.**
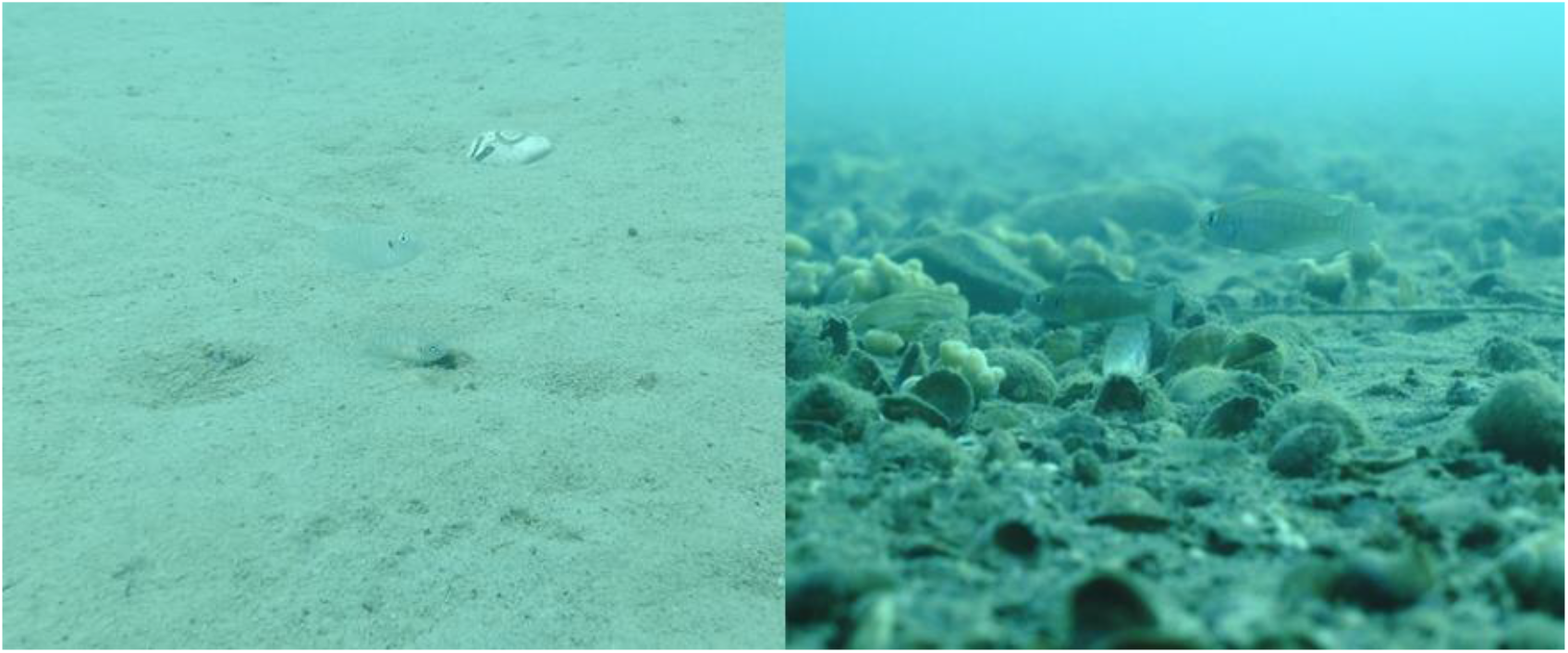
*N. brevis population from separated shells on sand (****S***, left); right, *N. brevis population from shell bed (****B***, right).

*Neolamprologus brevis* from Chezi (8°46’44.9”S, 31°00’25.2”E) reside in a sparse environment of separated shells (hence we refer to this population as *S* for “separated shells”(Ota et al., 2012)), and form breeding pairs inhabiting individual shells scattered across open sandy habitats. Pairs maintain small territories, engaging in low-frequency social interactions. Females are smaller than males and both sexes inhabit in the same shell, which serves as a brooding site and is actively defended from intruders.

The Chikonde Island population (8°42′43″S, 31°05′33″E) reside in dense shell beds (population hereafter ***B*** for “shell bed”, (Ota et al., 2012)). These individuals are similar in size, preventing them from cohabitating in a single shell and territories within colonies are not visually distinct (BM and AJ pers obs.). Both locales share similar diets, feeding primarily on invertebrates via plankton snatching and substrate feeding (Konings, 2019), and experience comparable predation pressures from *Lepidiolamprologus elongatus*, a widespread piscivore (Lein & Jordan, 2021).

There has been some argument for returning individual species status of these two forms (that ***B*** be reinstated to *Neolamprologus calliurus*), with meaningful evidence both for and against e.g. (Koblmüller et al., 2007), we defer to the findings of (Ota et al., 2012) which represent the most recent addition to this discussion, the results of which concur greatly with our own, thus refer to the two populations as separate ecomorphs of *N. brevis* hereafter.

### 2. Behavioural Observations

Behavioural data were collected from eight ***S*** groups at Chezi and nine ***B*** at Chikonde to assess social interactions and foraging activity. Observations were conducted between April and May 2022 at Chezi and in May 2023 at Chikonde Island. Each group was observed for 30–50 minutes using SCUBA diving and underwater video recordings (GoPro Hero6). behavioural recording occurred between 10:00 AM and 11:00 AM to standardize diurnal activity patterns. Videos were analysed using BORIS software (Friard & Gamba, 2016). Behaviour scoring was conducted on a single focal individual randomly chosen in the group. The first and last 10 minutes of each recording were excluded to minimize disturbance effects during diver entry and camera retrieval. Social interactions (e.g., courtship, aggression, and affiliative behaviours) were classified based on an ethogram adapted from (Reyes-Contreras et al., 2019). Foraging behaviour (e.g., substrate and plankton feeding) was recorded as a proxy for energy intake. Behaviour was quantified as frequencies by dividing the total number of each behaviour exhibited by the observation time (in minutes).

### 3. Sampling and Brain Morphology Measurements

Following behavioural observations, two individuals from each filmed group were captured and euthanized using an overdose of MS-222 (tricaine methanesulfonate, 1 g/L). Standard length and body weight were measured. Fish heads were fixed in a solution of 4% paraformaldehyde and 1% glutaraldehyde in phosphate buffer at 4°C for 24 hours. Brains were then extracted and preserved in fresh fixative for one week before measurements. Brains were photographed from dorsal, ventral, and lateral perspectives using a stereo zoom microscope (AXIO Zoom.V16, ZEISS) equipped with a digital camera (Axiocam 503 mono, ZEISS). Five brain regions—the telencephalon, optic tectum, cerebellum, dorsal medulla, and hypothalamus—were selected based on their functional relevance to social and foraging behaviours (Pollen et al., 2007). Each region’s length (L), width (W), and height (H) were measured using ImageJ software (Schneider et al., 2012). Volumes were calculated using the ellipsoid formula: V=6π×L×W×H.

Only adult individuals were included in the analysis, and sex was confirmed via gonadal examination post-dissection. Additional brain samples (that were not recorded for behaviour) from eight ***S*** individuals and one ***B*** were included in the study. In total, 43 individuals were analyzed (***S***: 24; ***B***: 19).

### 4. Data Analysis

Behavioural frequencies were analysed to compare social and foraging activity across populations. Separate linear models (LMs) were used to examine differences in the frequency of social interactions and foraging behaviours, with behavioural frequency as the response variable and population as the predictor. For total brain volume, linear models were compared using Akaike Information Criterion (AIC) to select the best-fit model. The individual’s sex did not significantly influence the brain volume after accounting for the body weight. The final model included an interaction term between population and log-transformed body weight, accounting the differences in the allometric relationship between body weight and brain volume across populations. Brain region analyses were conducted using separate linear models for each of the five regions. The response variable was the log-transformed volume of the brain region, and the predictors included population and the log-transformed volume of the rest of the brain (total brain volume minus the region of interest). All statistical analyses were conducted in R (v4.2.0). Residual diagnostics were performed to confirm model assumptions of normality and homoscedasticity. Statistical significance was set at p<0.05.

### 5. Ethical Considerations

All methods adhered to the ASAB/ABS Guidelines for the Use of Animals in Research. Field work was carried out with approval from the Fisheries Department at the Ministry of Fisheries and Livestock Zambia, under study permits issued by the government of Zambia (No. G7067690 and C3195368, SP260718/7-21, SP425444/4-24) and in conjunction with a memorandum of understanding with the University of Zambia (MOU 101/14/11). The study species is listed as ‘Least Concern’ on the IUCN Red List of Threatened Species.

## Results

### 1. behavioural Differences Between Populations

We quantified social and foraging behaviours in the ***S*** and ***B*** populations (**Figure 2**). The frequency of social interactions was **significantly higher** in ***B*** compared to ***S*** population (0.83 ± 0.06 vs. 0.22 ± 0.09; β = 0.612, p < 0.001). In contrast, the frequency of foraging behaviour (food pecking) **was not significantly different** between the two populations (***B***: 0.55 ± 0.07 vs. ***S***: 0.58 ± 0.08; β = -0.023, p = 0.835).

**Figure 2.**
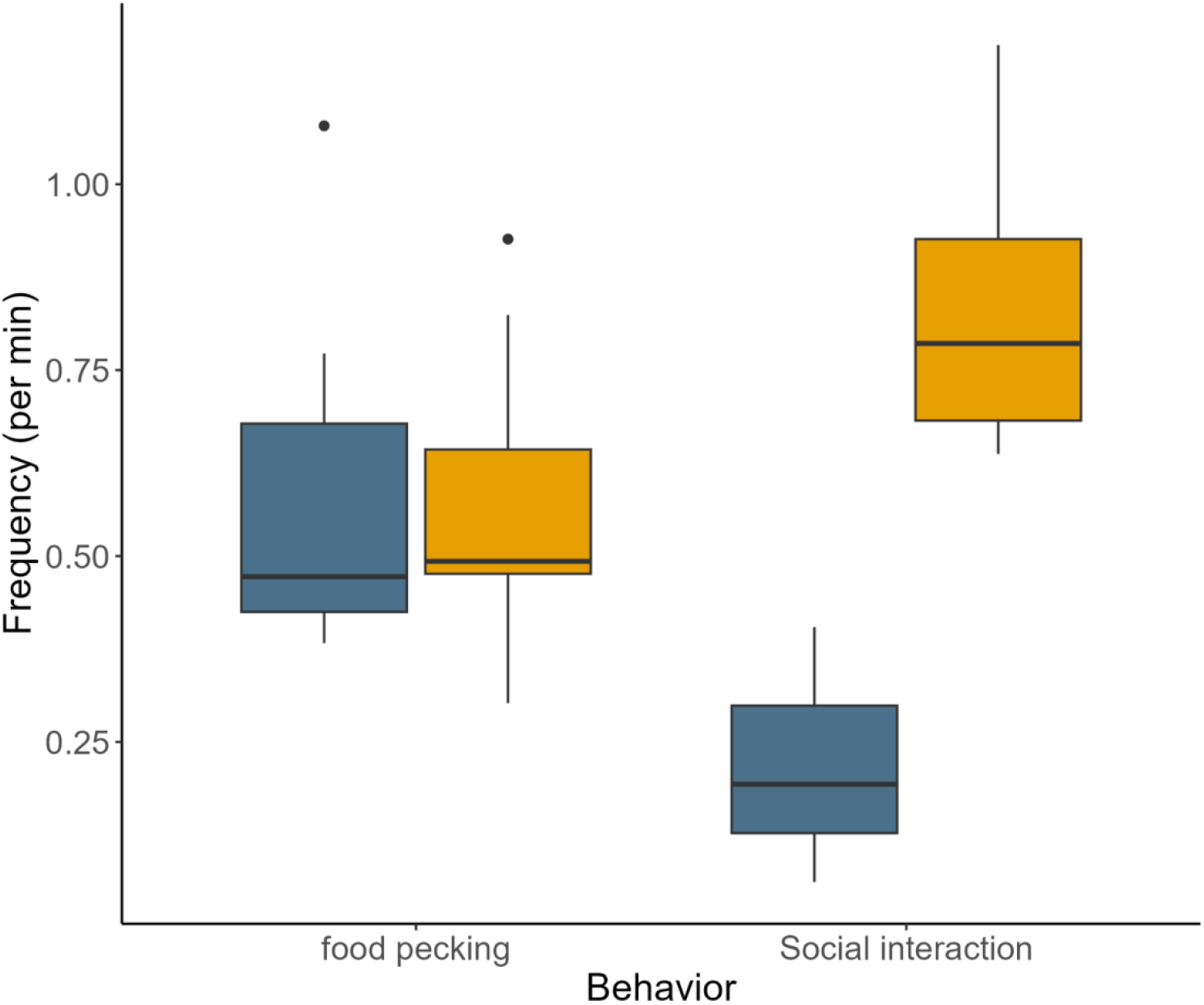
Comparison of behavior frequencies (per minute) between sand and shells (“*S*” in text, Chikonde, blue bars) shell bed (“*B*” in text, Chezi, yellow bars) populations. The two behavioural categories analysed were “feeding” and “social interaction.” Populations on shell beds (***B***) fish exhibited a higher frequency of social interactions than those on sand ans shells (***S***), while both populations showed similar frequencies of feeding.

### 2. Total Brain Volume and Body Weight

The relationship between total brain volume and body weight differed significantly between populations (**Figure 3a**). The model revealed a significant interaction between population and log-transformed body weight (β = 0.261, p = 0.015), indicating that the effect of body weight on brain volume varied between populations. Specifically, after controlling for body weight, ***B*** had a **significantly greater** brain volume compared to ***S*** (β = 0.321, p < 0.001). We also examined the relationship between body weight and standard length (**Figure 3b**). After accounting for standard length, ***S*** exhibited significantly higher body weights than ***B*** (β = 0.171, p = 0.013).

**Figure 3.**
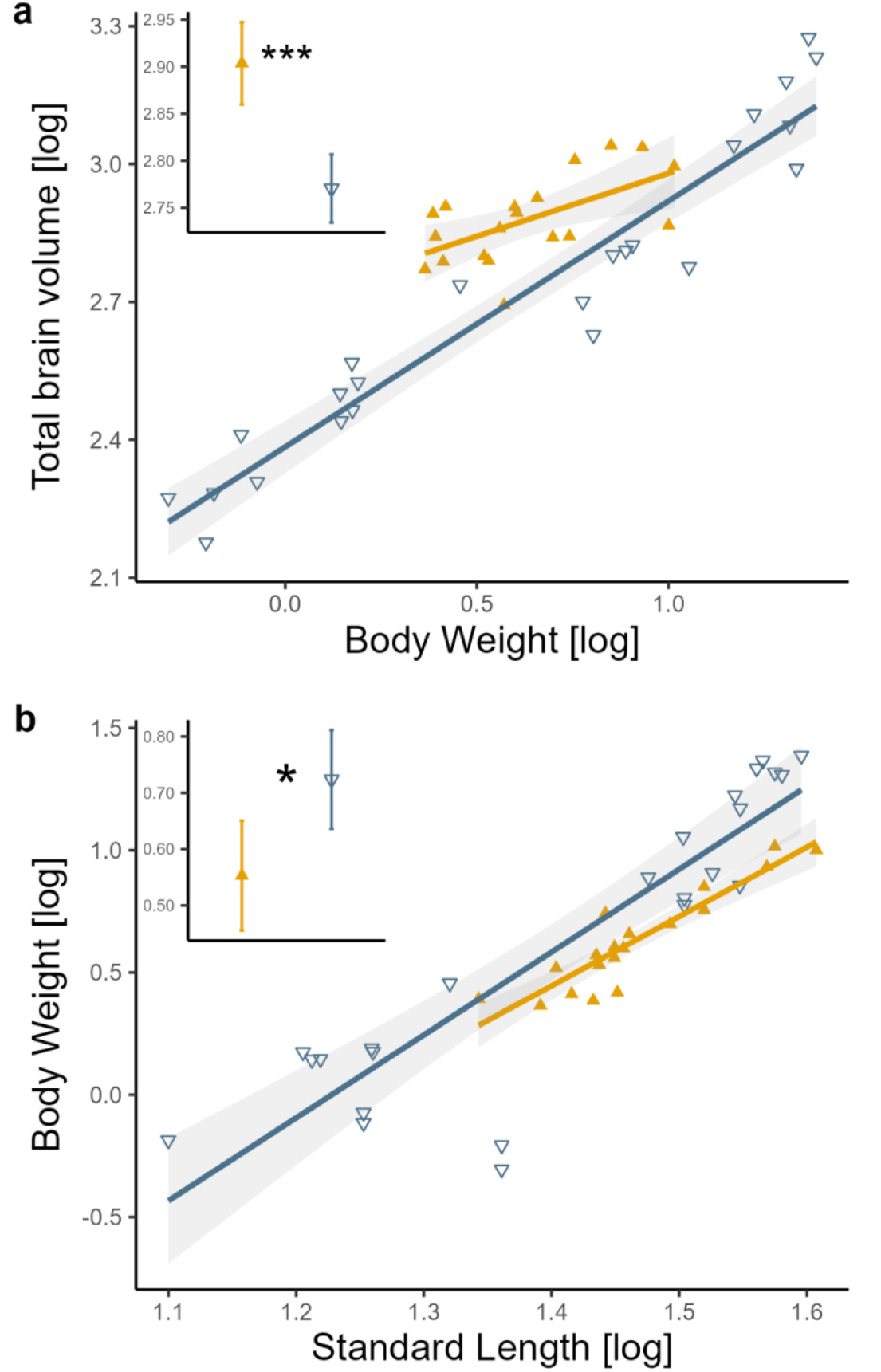
Relationship between total brain volume, body weight, and standard length in shell bed (“B” in text, Chikonde, blue points) and sand and shells (“S” in text, Chezi, yellow points) populations. (a) Relationship between total brain volume (log-transformed) and body weight (log-transformed). (b) Relationship between body weight (log-transformed) and standard length (log-transformed). The shaded regions represent 95% confidence intervals, and the insets show predicted values with the corresponding confidence intervals. Asterisks (***) and (*) denote statistically significant differences (p < 0.001 and p < 0.05, respectively).

### 3. Relative Brain Region Volumes

Relative brain region volumes were analysed to investigate potential brain structure differences between populations (**Figure 4**). Compared to ***S***, the ***B*** population exhibited a **significantly larger** relative telencephalon volume (β = 0.109, p = 0.021) and a **smaller** relative hypothalamus volume (β = -0.110, p = 0.034). No significant differences were observed for the optic tectum (β = 0.017, p = 0.635), cerebellum (β = -0.007, p = 0.868), or dorsal medulla (β = -0.072, p = 0.401).

**Figure 4.**
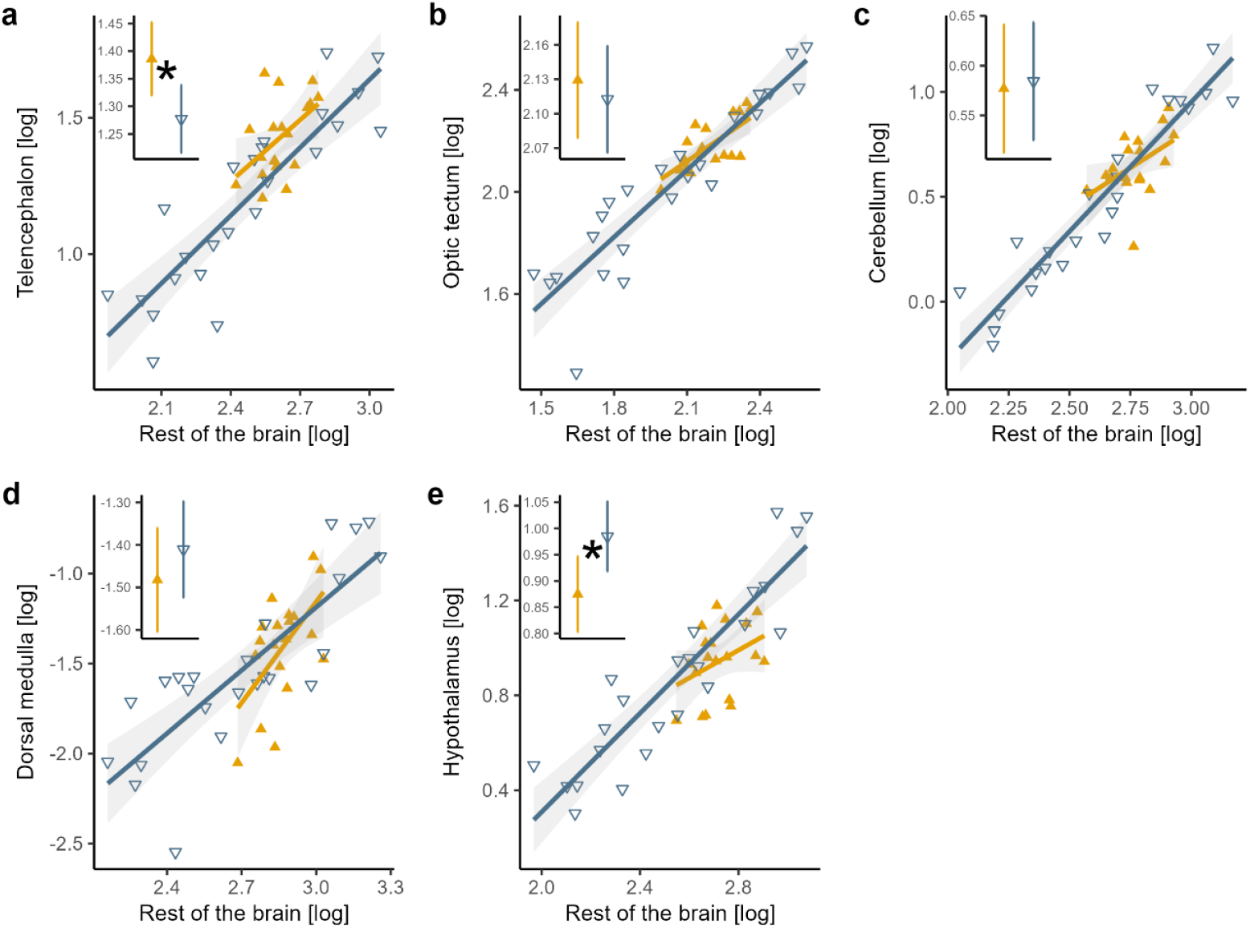
Relationship between individual brain regions and the rest of the brain in the shell bed (“B” in text, Chikonde, blue points) and sand and shells (“S” in text, Chezi, yellow points) populations. (a) Telencephalon, (b) Optic tectum, (c) Cerebellum, (d) Dorsal medulla, and (e) Hypothalamus. All volumes were log-transformed. The shaded regions represent 95% confidence intervals, and the insets show predicted brain region volumes with corresponding confidence intervals. Asterisks (*) denote statistically significant differences (p < 0.05).

## Discussion

In this study, we leveraged variation in social complexity across populations in the same species to test the predictions of the social brain hypothesis in natural populations. Specifically, the population on dense shell beds (***B***) exhibited more frequent social interactions and larger brain volumes relative to body size than the population on sparse sand and shells (***S***), consistent with our predictions. Furthermore, the telencephalon was relatively larger in ***B***, while the hypothalamus was smaller. These results suggest that enhanced cognitive demands in complex social environments may drive both overall brain enlargement and regionspecific adaptations.

Previous research on neuroanatomical traits and social complexity has produced mixed results, and our work suggests this may in part be due to methodological challenges and ecological confounds. Our results are consistent with findings from some interspecific studies that link social complexity to change in specific brain regions. For example, Sakai et al. (2011) demonstrated that the most socially complex hyena species, possess the largest relative brain sizes and frontal cortex volumes. Similarly, in cichlid fish, Pollen et al. (2007) observed larger telencephalons in monogamous cichlid species. These findings collectively support the hypothesis that increased social demands drive neuroanatomical specializations, particularly in brain regions associated with social cognition. However, interspecific studies in birds and rodents show the opposite relationship between brain size and social complexity. For instance, Hardie and Cooney (2023) found that developmental mode and foraging niche were stronger predictors of brain size than social factors in 1886 bird species. Matějů et al. (2016) reported that while sociality increased absolute brain size in ground squirrels, relative brain size did not correlate with social complexity. Similarly, in African mole-rats, Kverková (2018) observed larger absolute brain sizes and forebrain neuron counts in solitary species compared to social ones. Finally, Benson-Amram et al. (2016) found that relative brain size predicted problem-solving abilities in carnivores but noted no relationship between social complexity and brain size. These contrasting findings highlight the intricate relationship between sociality and brain evolution, and underscore the challenges posed by ecological confounding factors inherent in interspecies comparisons, as well as the methodological difficulties encountered in studies involving wild animals. Our work, specifically testing within species that naturally differ in their social organization, allows us to quantify the impact of social environment and development on brain morphology without some of these confounds of inter-species comparisons. We conducted our research in the field with animals of known providence, leveraging standing variation in social systems among populations. A similar intraspecies approach previously found that cleaner wrasse living in high-density social environments had larger forebrains (Triki et al., 2019). This type of field-based approach, using existing variation in social behaviour, minimizes confounding factors and provides a robust framework for testing hypotheses about the role of sociality in shaping brain size and structure.

We use direct behavioural observations to provide a comprehensive perspective on social complexity, avoiding broad categories (e.g. low vs high complexity) or proxies of social interaction (e.g. group size, animal density, or mating system). The increased frequency of social behaviour in the shell bed population (***B***) may be a response to a more active local conspecific environment, with frequent engagement with neighbours who are encountered more often due to their higher density. Maintaining social relationships with many neighbours is thought to be energetically and cognitively costly (Dunbar, 2018) and we found that the shell bed population, which had a four-fold higher interaction rate than the sand and shell population, had concomitantly larger brains. Such a finding was only possible through the use of direct behavioural observation of wild animals in combination with measurements of neuroanatomy.

This approach also allows us to assess whether energy intake plays a role in shaping the brain in our population comparison. Brain tissue is metabolically expensive to maintain, imposing strong selective pressures against nonadaptive changes (A. Kotrschal et al., 2013; Leslie C. Aiello et al., 1995; Navarrete et al., 2011). In our study, the similar frequency of foraging behaviour, as well as shared foraging niche among populations suggests energy intake was unlikely to differ across populations. Indeed, the population with the highest body condition (sand and shells) had smaller relative brain volumes, providing strong evidence that higher energy intakes were not a major causative factor for differences in the brain. Rather, the major difference appears to be due to the increased social interactions, an interpretation supported by the observed enlargement in the shell bed population ***B*** of the telencephalon, a region known to mediate complex social behaviours, such as decision-making (O’Connell & Hofmann, 2012), and social cognition (Triki et al., 2022). In contrast, the reduced hypothalamus in ***B*** may appear counter-intuitive, as this region also plays a role in maintaining sociality, with cichlids raised in denser social groups demonstrating an enlarged hypothalamus (Fischer et al., 2015), as well as regulating endocrine pathways involved with (predominantly affiliative) social behaviours in a reproductive context (Demski & Knigge, 1971). As such the reduction in hypothalamus volume may instead reflect a trade-off, a consequence of the mosaic brain hypothesis (Barton & Harvey, 2000), which suggests a zero-sum interpretation of brain development. In this scenario, energy and resources may have been prioritized in brain regions supporting social and cognitive functions in the more socially complex shell bed population.

Although our study provides evidence of a positive relationship between social complexity and brain size, it does not establish causality. Is increased sociality experienced by an organism driving the expansion of brain size, or are organisms with larger brains inherently more socially competent, enabling them to thrive in more socially complex environments? Studies tracking neuroanatomical changes in response to experimentally manipulated social environments are needed to disentangle these effects. For example, laboratory experiments where social stimuli can be systematically varied, could provide more direct evidence for causal relationships. Additionally, while our intraspecies comparisons can reduce the potential ecological confounds, these two populations are still different in their genetics and location. Comparing individuals within a single population that vary in social complexity would further remove these confounds, enabling a more nuanced resolution of the social effect on the role of sociality in brain size variation.

Overall, our study demonstrates that increased social complexity is associated with larger brain size and specific neuroanatomical adaptations in cichlid *N. brevis*, providing compelling empirical support for the Social Brain Hypothesis. Moreover, our findings emphasize the importance of integrating behavioural and neuroanatomical data to validate the social effect on brain anatomy. Finally, our research underscores the value of cichlid fish as a model for studying brain evolution and highlights the potential of using intraspecific design to explore the intricate interplay between sociality and brain morphology.

## Acknowledgements

We thank the Department of Fisheries in Mpulungu, Zambia for kindly supporting our research at Lake Tanganyika. We are grateful to the staff at the Tanganyika Science Lodge and Kalambo Falls Lodge for their hospitality.

## Contribution Statement

**Bin Ma:** Conceptualization (lead); data collection and curation (lead); formal analysis (lead); investigation (lead); methodology (lead); project administration (equal); resources (equal); visualization (lead); writing – original draft (lead); writing – review and editing (lead). **Boyd Dunster**: data collection (lead); writing – review and editing (supporting); resources (supporting). **Etienne Lein:** data collection (lead); resources (supporting). **Boshan Zhu**: data collection (supporting). **Weiwei Li:** data collection (supporting); **Alex Jordan**: Supervision (lead); Funding acquisition (lead); project administration (lead); resources (lead); writing – review and editing (co-lead).

## Data Accessibility

All associated data are available at DRYAD

## References

Barton, R. A., & Harvey, P. H. (2000). Mosaic evolution of brain structure in mammals. Nature, 405(6790), 1055–1058. 10.1038/35016580

Benson-Amram, S., Dantzer, B., Stricker, G., Swanson, E. M., & Holekamp, K. E. (2016). Brain size predicts problem-solving ability in mammalian carnivores. Proceedings of the National Academy of Sciences of the United States of America, 113(9), 2532–2537. 10.1073/pnas.1505913113

Demski, L. S., & Knigge, K. M. (1971). The telencephalon and hypothalamus of the bluegill (Lepomis macrochirus): Evoked feeding, aggressive and reproductive behavior with representative frontal sections. Journal of Comparative Neurology, 143(1), 1–16. 10.1002/cne.901430102

Dunbar, R. I. M. (2009). The social brain hypothesis and its implications for social evolution. Annals of Human Biology, 36(5), 562–572. 10.1080/03014460902960289

Dunbar, R. I. M. (2018). The Anatomy of Friendship. Trends in Cognitive Sciences, 22(1), 32–51. 10.1016/j.tics.2017.10.004

Finarelli, J. A., & Flynn, J. J. (2009). Brain-size evolution and sociality in Carnivora. Proceedings of the National Academy of Sciences, 106(23), 9345–9349. 10.1073/pnas.0901780106

Fischer, S., Bessert-Nettelbeck, M., Kotrschal, A., & Taborsky, B. (2015). Rearing-Group Size Determines Social Competence and Brain Structure in a Cooperatively Breeding Cichlid. The American Naturalist, 186(1), 123–140. 10.1086/681636

Friard, O., & Gamba, M. (2016). BORIS: A free, versatile open-source event-logging software for video/audio coding and live observations. Methods in Ecology and Evolution, 7(11), 1325–1330. 10.1111/2041-210X.12584

Hardie, J. L., & Cooney, C. R. (2023). Sociality, ecology and developmental constraints predict variation in brain size across birds. Journal of Evolutionary Biology, 36(1), 144–155.

Koblmüller, S., Duftner, N., Sefc, K. M., Aibara, M., Stipacek, M., Blanc, M., Egger, B., & Sturmbauer, C. (2007). Reticulate phylogeny of gastropod-shell-breeding cichlids from Lake Tanganyika – the result of repeated introgressive hybridization. BMC Evolutionary Biology, 7(1), 7. 10.1186/1471-2148-7-7

Konings, A. (2019). Tanganyika Cichlids in Their Natural Habitat. Cichlid Press.

Kotrschal, A., Rogell, B., Bundsen, A., Svensson, B., Zajitschek, S., Brännström, I., Immler, S., Maklakov, A. A., & Kolm, N. (2013). Artificial selection on relative brain size in the guppy reveals costs and benefits of evolving a larger brain. Current Biology, 23(2), 168–171. Scopus. 10.1016/j.cub.2012.11.058

Kotrschal, K., Van Staaden, M., & Huber, R. (1998). Fish brains: Evolution and environmental relationships. REVIEWS IN FISH BIOLOGY AND FISHERIES, 8(4), 373–408. 10.1023/A:1008839605380

Kverková, K. (2018). Sociality does not drive the evolution of large brains in eusocial African mole rats.

Lein, E., & Jordan, A. (2021). Studying the evolution of social behaviour in one of Darwin’s Dreamponds: A case for the Lamprologine shell-dwelling cichlids. Hydrobiologia, 848(16), 3699–3726. 10.1007/s10750-020-04473-x

Leslie C. Aiello, Leslie C. Aiello, Lloyd Paul Aiello, P.E. Wheeler, & P.E. Wheeler. (1995). The Expensive-Tissue Hypothesis: The Brain and the Digestive System in Human and Primate Evolution. Current Anthropology. 10.1086/204350

Matějů, J., Kratochvíl, L., Pavelková, Z., Pavelková Řičánková, V., Vohralík, V., & Němec, P. (2016). Absolute, not relative brain size correlates with sociality in ground squirrels. Proceedings of the Royal Society B: Biological Sciences, 283(1827), 20152725. 10.1098/rspb.2015.2725

Navarrete, A., van Schaik, C. P., & Isler, K. (2011). Energetics and the evolution of human brain size. Nature, 480(7375), 91–93. 10.1038/nature10629

O’Connell, L. A., & Hofmann, H. A. (2012). Evolution of a Vertebrate Social Decision-Making Network. Science, 336(6085), 1154–1157. 10.1126/science.1218889

Ota, K., Aibara, M., Morita, M., Awata, S., Hori, M., & Kohda, M. (2012). Alternative Reproductive Tactics in the Shell-Brooding Lake Tanganyika Cichlid Neolamprologus brevis. International Journal of Evolutionary Biology, 2012, e193235. 10.1155/2012/193235

Pollen, A. A., Dobberfuhl, A. P., Scace, J., Igulu, M. M., Renn, S. C. P., Shumway, C. A., & Hofmann, H. A. (2007). Environmental Complexity and Social Organization Sculpt the Brain in Lake Tanganyikan Cichlid Fish. Brain, Behavior and Evolution, 70(1), 21–39. 10.1159/000101067

Reyes-Contreras, M., Glauser, G., Rennison, D. J., & Taborsky, B. (2019). Early-life manipulation of cortisol and its receptor alters stress axis programming and social competence. Philosophical Transactions of the Royal Society B: Biological Sciences, 374(1770), 20180119. 10.1098/rstb.2018.0119

Sakai, S. T., Arsznov, B. M., Lundrigan, B. L., & Holekamp, K. E. (2011). Virtual endocasts: An application of computed tomography in the study of brain variation among hyenas. Annals of the New York Academy of Sciences, 1225(SUPPL. 1), E160–E170. Scopus. 10.1111/j.1749-6632.2011.05988.x

Schneider, C. A., Rasband, W. S., & Eliceiri, K. W. (2012). NIH Image to ImageJ: 25 years of image analysis. Nature Methods, 9(7), 671–675. 10.1038/nmeth.2089

Triki, Z., Granell-Ruiz, M., Fong, S., Amcoff, M., & Kolm, N. (2022). Brain morphology correlates of learning and cognitive flexibility in a fish species (Poecilia reticulata). Proceedings of the Royal Society B: Biological Sciences, 289(1978), 20220844. 10.1098/rspb.2022.0844

Triki, Z., Levorato, E., McNeely, W., Marshall, J., & Bshary, R. (2019). Population densities predict forebrain size variation in the cleaner fish Labroides dimidiatus. Proceedings of the Royal Society B: Biological Sciences, 286(1915), 20192108. 10.1098/rspb.2019.2108

